# Humic acids enrich the plant microbiota with bacterial candidates for the suppression of pathogens

**DOI:** 10.1101/2020.07.18.210427

**Authors:** Maura Santos Reis de Andrade da Silva, Orlando Carlos Huertas Tavares, Thiago Gonçalves Ribeiro, Camilla Santos Reis de Andrade da Silva, Carolina Santos Reis de Andrade da Silva, José Maria García-Mina, Vera Lúcia Divan Baldani, Andrés Calderín García, Ricardo Luiz Louro Berbara, Ederson da Conceição Jesus

**Affiliations:** Embrapa Agrobiologia – Rodovia BR 465, Km 7, Bairro Ecologia, CEP 23891-000, Seropédica, RJ/Brasil; Universidade Federal Rural do Rio de Janeiro, RJ/Brasil; Universidade de Navarra, Espanha

**Keywords:** humic substances, metaxonomics, plant microbiome, plant-growth promotion, plant-microbe interactions, plant defense

## Abstract

Humic acids (HAs) stimulate the growth of several plant species by regulating their hormonal and redox metabolisms. Nevertheless, studies on the relationship of these substances with the plant-associated microbiota are almost nonexistent. Here, we hypothesized that the effect of HAs occurs in parallel with the regulation of the plant-associated bacterial community. Our results show the positive influence of HAs on the growth of rice and its stimulation of the root system. Metataxonomics revealed that the structure and composition of root bacterial communities were affected upon the application of HAs. *Chitinophaga* and *Mucilaginibacter* were the predominant genera in HA-treated roots. These bacteria produce enzymes that degrade compounds like those present in the wall of fungi, oomycetes, and nematode eggs. *Pseudomonas* and the *Gp 1* group of *Acidobacteria,* both siderophore-producers and plant-growth promoters were also enriched, although with lower abundances. Given these results, we suggest that plants recruit these microorganisms in response to the stress caused by the HA-root interaction. For the first time, our findings indicate that HA-stimulated plants adopt the ecological strategy of recruiting members of the bacterial community that are candidates for the suppression of pathogens and, therefore, involved in plant defense.

## Introduction

Humic substances (HS) benefit plants through their effects on soil physical, chemical, and biological properties. These substances can also promote plant growth, including changes in root thickness, length, branching, and density, thus altering root architecture (Canellas and Olivares, 2014; Garcia *et al*., 2016 a*b*; Tavares *et al*., 2020). Nevertheless, its effects are not restricted to morphological changes. Humic acids (HA) regulate hormonal signaling pathways. Some scientists suggest that humic fragments can be mimic auxins, being recognized as such by cellular receptors (Canellas *et al.*, 2002). Olaetxea *et al*. (2015) showed that HA regulates the gene expression of membrane aquaporins (PIP) under stress conditions through ABA’s pathways. These substances also regulate the plant redox metabolism, modifying the concentration of reactive oxygen species (ROS) and stimulating the expression of scavenger peroxidases, -POX, CAT-catalases, and tonoplast aquaporins (TIPs) (Garcia *et al*., 2016*a*).

By affecting the plant metabolism, HS can also alter the root exudation profile, thus interfering with the structure of the rhizosphere microbial community (Puglisi *et al*., 2009; Puglisi *et al*., 2013). HS structural complexity makes these substances elicit several metabolic pathways and, consequently, interact with plant-associated microorganisms. Several studies report the benefits of applying both HS and growth-promoting bacteria exogenously (Marques Júnior *et al*., 2008; Canellas *et al*., 2013; Olivares *et al*., 2015). As an example, we can cite the differential biofilm production and colonization of corn roots by *Herbaspirillum seropedicae* upon the application of HA (Canellas and Olivares, 2017), and the protection of plants against water stress adverse effects when co-inoculated with bacteria and HA (Aguiar *et al*., 2016). HA stimulate the activity of oxidative metabolic enzymes (SOD, CAT, and APX); in turn, bacteria induce the preservation of relative water content and stomata (Aguiar *et al.*, 2016).

It is well established in the literature that plants naturally harbor a high microbial diversity in their tissues. This microbiota affects plant growth and survival (Vandenkoornhuyse *et al*., 2015) and is highly responsive to changes in plant physiology (Hallmann and Berg, 2006). Despite its importance, studies on the effects of HA on the plant-associated microbial community are almost nonexistent. In this scenario, we hypothesize that the effect of HA occurs in parallel with the regulation of the plant-associated bacterial community responsible for promoting plant growth and protecting plants from biotic and abiotic stresses. To test this hypothesis, we applied humic acids (HA), the HS fraction soluble in basic pH, precipitates in an acid medium (Schnitzer, 1978), on rice seedling roots. HA were added to the nutrient solution every 72 hours for 264 hours. We collected plants successively every 24 hours to assess their development. The quantitative growth and root morphological variables, together with the taxonomic characterization of the bacterial community by sequencing 16S rRNA genes, were evaluated.

## Material and Methods

### Extraction, purification, and characterization of humic acids

Humic acids (HAs) were isolated from bovine manure vermicompost and purified according to the methodology recommended by the International Humic Substances Society (IHSS, 2016). Their characteristics were previously described (Tavares *et al*., 2020).

### Experimental design, sampling, and evaluation of the morphological parameters of rice plant roots

The experiment was carried out in a completely randomized design. The treatments consisted of combining the levels of two factors, HA application (with and without HA) and sampling time (11 sampling times every 24 h throughout the experiment), totaling 22 treatments, with six replicates each.

Rice plants of the Piauí variety were grown in a growth chamber under a 12 h/12 h (light/dark) photoperiod regime with 480 μmol m^−2^ s^−1^ of photosynthetic photon flow, 70% relative humidity, and temperatures of 28°C/24°C (day/night). The seeds were disinfected in 2% sodium hypochlorite for 30 minutes in an orbital shaking and then washed ten times in distilled water. Seven days after germination, the seedlings were transferred to pots with a capacity of 0.7 L (four plants per pot) in Hoagland’s solution (Hoagland and Arnon, 1950).

The treatments consisted of the application or not of HAs at a dose of 80 mg HA L^−1^in nutrient solution. HAs were added to the nutrient solution every 72 hours when the nutrient solution was changed. Plants were collected every 24 h until 264 h (11 days) after transplantation, and the following variables were measured: root and shoot dry mass; leaf area; root length, surface area, and volume; and the number of root tips. Leaf area was estimated with a photoelectric meter (LI-3000, Li-Cor). The roots were scanned with an Epson Expression 10000XL scanning system with an additional light unit (Turbo Pascal Unit, TPU). Root length (mm), surface area (mm^2^), volume (mm^3^), and the number of root tips were analyzed with the WinRhizo Arabidopsis software, 2012b (Régent Instruments Inc., Quebec, Canada).

Data normality and homoscedasticity were tested with the Shapiro-Wilk and Bartlett’s tests, respectively. The data were submitted to analysis of variance (ANOVA) (p < 0.05) using the R software (R Development Core Team, 2017).

### Quantitative growth analysis

Plant growth was determined as the total dry weight and leaf area index. The original data were adjusted by non-linear regression, deriving the growth rates, and estimates of the rates’ instantaneous values. A logistics model based on iterative processes selected with basis on the significance of the model coefficients, determination coefficient (R^2^) and the global trend of temporal variation of the measured variables. The absolute (AGR) and relative growth (RGR) and net assimilation rates (NAR) were obtained from the logistics function for dry weight and leaf area index (Hunt, 1982).

### DNA extraction and 16S rRNA sequencing

Rice plants treated or not with HAs were collected 240 h after the first HA application and used for DNA extraction and sequencing of the 16S rRNA gene. Total DNA was extracted from shoot and roots using the PowerPlant® Pro DNA Isolation Kit (MO BIO) according to the manufacturer’s protocol. The measurement of DNA quality was performed by electrophoresis on 1% agarose gel and visualization through gel staining with ethidium bromide. DNA quantification was carried out by spectrophotometry in NanoDrop 1000. The V4 region of the 16S rRNA gene was sequenced with the 515F (GTGCCAGCMGCCGCGGTAA) and 806R (GGACTACHVGGGTWTCTAAT) primers as previously described Caporaso et al. (2012) and following the Earth Microbiome Project’s protocol (http://www.earthmicrobiome.org/protocols-and-standards/16s/). Peptide nucleic acid (PNA) clamps were used to block the amplification of chloroplast and mitochondrial sequences (Fitzpatrick et al., 2018). Sequencing was carried out with a MiSeq sequencing machine at the Environmental Sample Preparation and Sequencing Facility of the Argonne National Laboratory.

### Sequence analysis

The libraries were demultiplexed with CASAVA v.1.8 (Illumina) and deposited at Genbank’s Sequence Read Archive under the numbers SAMN5464651 through SAMN5464666 (Supplementary Table 2). The sequences were analyzed using the open-source software Mothur (v.1.41.1) (Schloss et al., 2009) and following the MiSeq SOP guidelines (https://www.mothur.org/wiki/MiSeq_SOP). Forward and reverse sequences were assembled, producing 288,780 total reads ranging between 247 and 502 base pairs with the average library size of 16,043 reads. Ambiguous sequences, sequences with more than 257 bp, and those with more than eight homopolymers were removed. A total of 245,783 sequences remained after these steps. Those were divided into 29,572 unique sequences, which were aligned with the V4 region of the reference database (SILVA bacterial release 132, http://www.arb-silva.de) (Quast et al., 2012; Yilmaz et al., 2014) using the standard settings for the global alignment method (Needleman and Wunsch, 1970). The aligned sequences were grouped with the command pre.cluster (diffs = 2) and checked for chimeras using the UCHIME software (Edgar et al., 2011), after which the chimeras were removed. The classification of the operational taxonomic units (OTUs) was performed with the Ribosomal Database Project Classifier tool (Cole et al., 2009) with an 80% bootstrap. Unknown OTUs and those that were classified as Eukaryota, Archaea, Mitochondria, and chloroplast were removed. The sequences were clustered using the average neighbor algorithm at 97% similarity (cutoff = 0.03), allowing the sequences to be classified at the genus level (Comeau et al., 2011).

The files were imported to the R software (R Development Core Team, 2017) and analyzed with the packages phyloseq (McMurdie and Holmes, 2013), stringr (Wickham, 2019), vegan (Oksanen, et al., 2019), DESeq2 (Love et al., 2014), and ggplot2 (Wickham, 2016). Bacterial genera whose abundances were differentially enriched or decreased in HA-treated plants were identified with the differential gene expression analysis on the negative binomial distribution implemented in package DESeq2 (Love et al., 2014). DESeq2 estimates variance-mean dependence in count data and tests for differential abundance based on a model using the negative binomial distribution. This approach was initially developed to identify differentially expressed genes in RNA-seq assays. However, it has also been used to identify differences in count data from metagenomic analysis. Boxplots were built to show that the abundance of the taxa that were significantly affected. Bar plots were built to show variations in the abundances of phyla and genera. Taxa with an abundance of < 2% were not shown. A principal component analysis was carried out on the covariance matrix calculated with OTU data.

## Results

### Effect of HA on biomass production, root growth, and development of rice plants

The effect of HA on root and shoot biomass accumulation varied throughout the evaluation period. Root biomass decreased 24 hours upon the first HA application, with a subsequent recovery after 48 h. The application of HA increased root biomass at 144 and 216 h (p <0.001) (Figure 1a). Shoot biomass had a similar response, with a significant increase at 144 and 264 h (p <0.001) (Figure 1b). The net assimilation rate (NAR) increased upon the application of HA (Figure 1e), indicating a stimulus to components of photosynthetic metabolism. Consequently, the absolute (AGR) and relative (RGR) growth rates increased at 100 h and remained so until the end of the experiment (Figure 1c,d).

**Figure 1.**
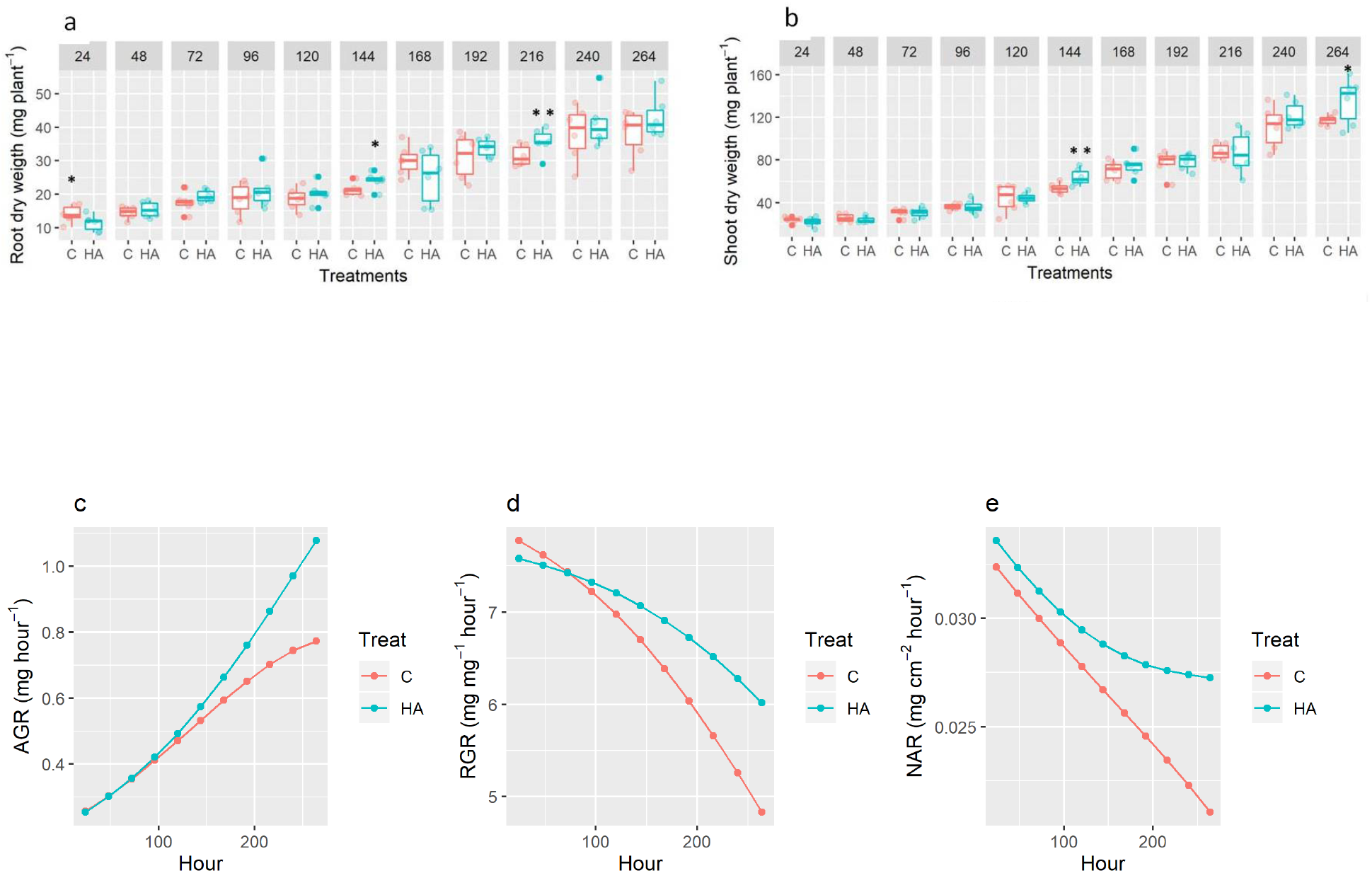
Root (a) and shoot (b) dry weight of rice plants treated (HA) or not (C) with 80 mg L^−1^ of humic acids in their roots. HA and C stand for humic acids and control, respectively. The asterisks indicate significant differences between treatments at each sampling time, as estimated by the F test (* for p < 0.10, and ** for p < 0.05). The numbers above the boxes show the hours after the first HA application. Variations in the net assimilation, absolute growth (d), and relative growth (e) rates after 264 hours.

HA stimulated significant increases in root biomass, length, surface area, and volume (Figures 1a and 2a,b,c). The moment when these effects were observed differed with the measured variable. Nevertheless, all of them pointed out to morphological changes and growth stimulus of the rice root system. The root volume showed a notable increase after 48 hours and throughout the remaining period of the experiment (Figure 2c). The effect of HA on root biomass was observed after 144 h (Figure 1a). Significant increases in root length and area were observed after 216 and 192 h, respectively (Figures 2a,b). The response of the number of tips was not significant (Figure 2d), except for an adverse response 144 h after the initial application (p <0.001).

**Figure 2.**
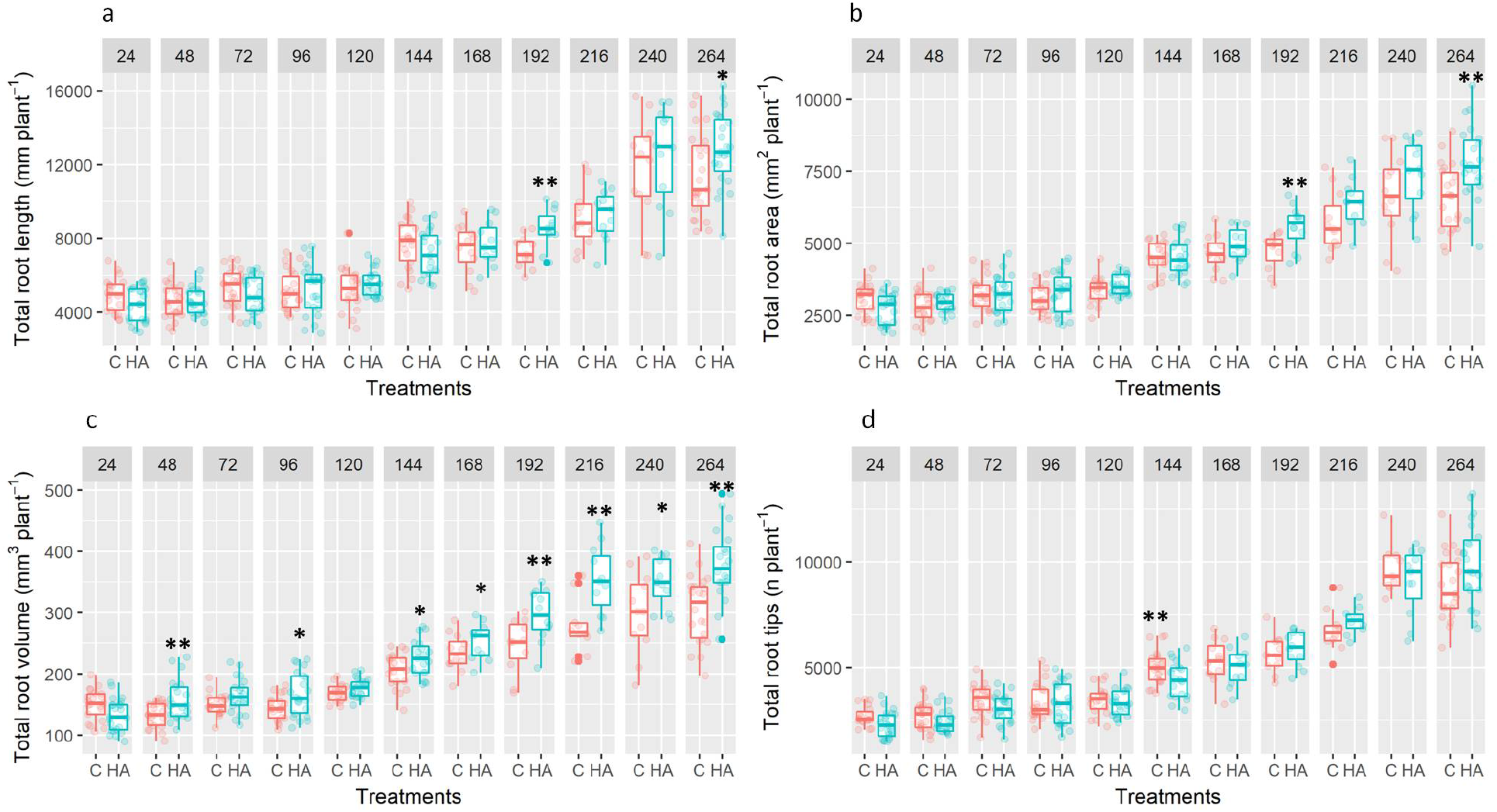
Root length (a), area (b), volume (c), and the number of tips (d) of rice plants treated (HA) or not (C) with 80 mg L^−1^ of humic acids in their roots. HA and C stand for humic acids and control, respectively. The asterisks indicate significant differences between treatments at each sampling time, as estimated by the F test (* for p < 0.10, and ** for p < 0.05). The numbers above the boxes show the hours after the first HA application.

### Changes in the community of bacteria associated with rice plants by the action of HA

Rice plants treated or not with HA were collected 240 h after the first HA application and used for DNA extraction and sequencing of the 16S rRNA gene. Using principal component analysis (PCA), we identified the division of the plant-associated bacterial communities into three major groups: one includes communities of roots treated with HAs; another contains communities of roots that did not receive these substances, and the other is formed by bacterial shoot communities (Fig. 3a). The variability in community composition was much higher in the roots than in the shoot. In fact, the shoot communities were very homogeneous concerning their taxonomic composition. Communities in the shoot were also more diverse, as shown by their higher number of phyla and genera (Fig. 3b,c).

**Figure 3.**
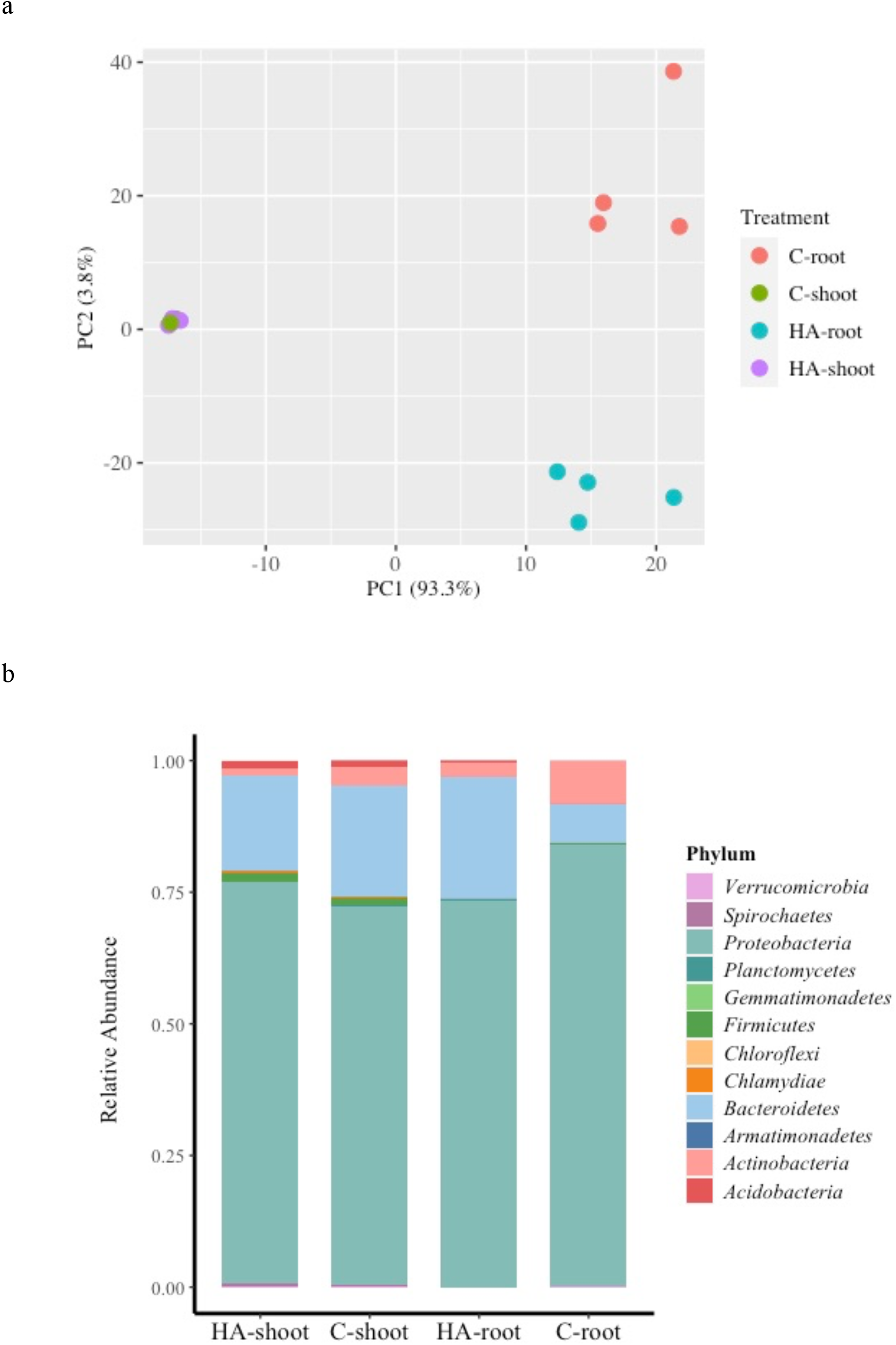
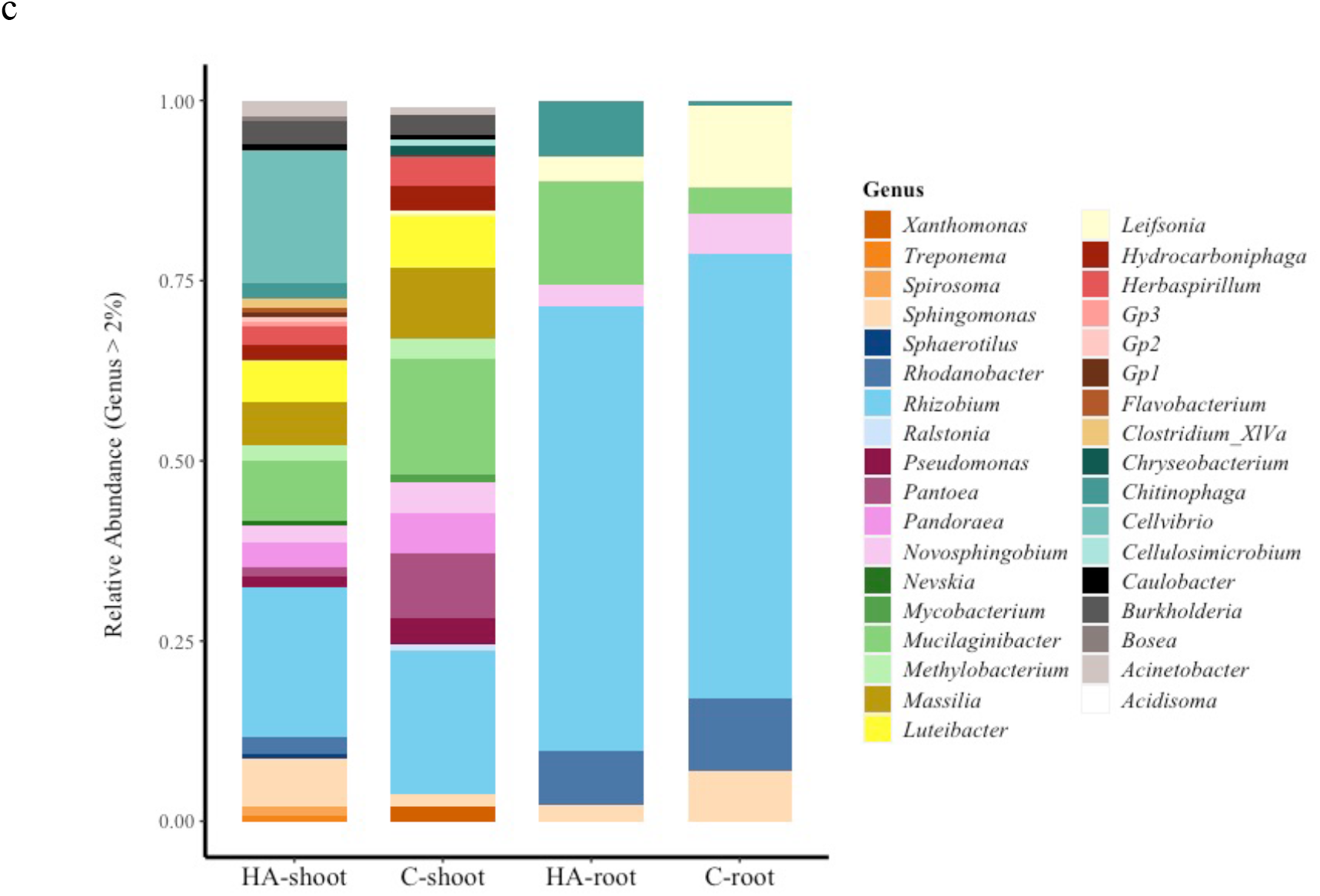
Bacterial communities differed between rice plants treated (HA) or not (C) with 80 mg L^−1^ of humic acids in their roots. (a) Principal component analysis showing the distance between the studied bacterial communities. C-shoot, C-root, HA- shoot, and HA-shoot stand for shoot and roots of the control and humic-acid-treated plants, respectively. The bar graphs display the relative abundance of taxa in roots and shoot classified at phylum (b) and gender (c) levels. In the case of genus, only those taxa with a relative abundance above 2% are displayed.

The metataxonomics approach made it possible to identify the microbial taxa that are modified by HA in plants in parallel with the stimulus they cause in plant growth. The bacterial community responded with an increase in the abundance of *Proteobacteria* and a decrease of *Bacteroidetes* in the shoots (Figure 3b). In turn, *Bacteroidetes* and *Acidobacteria* increased in the roots, and *Actinobacteria* decreased in both plant compartments. *Rhizobium* was the most abundant genus (Figure 3c), mainly in the roots, where it constituted more than 50% of the sequences. Contrary to our expectation, the abundances of bacteria which have been reported to promote plant growth, such as *Leifsonia*, *Sphingomonas,* and *Rhodanobacter* were reduced in HA-treated roots (Figure 3c and Figure4 a,b,c). In turn, the abundance of *Bacteroidetes* increased, especially those of the genera *Mucilaginibacter* and *Chitinophaga* (Figure 3 b,c and Figure 4d,e,f,g). This increase was statistically validated with DESeq analysis (Supplementary Table 1). *Pseudomonas* and a representative of *the Group 1 (Gp 1) of Acidobacteria* were also enriched, although with lower abundances (Figure 4h,i). Although the PCA has not shown a clear differentiation between shoot communities, we still can observe some differences in the abundance of specific genera. For example, the abundances of *Mucilaginibacter*, *Pantoea*, *Pandoraea*, *Pseudomonas*, and *Massilia* decreased in the shoot of plants treated with HA (Figure 4 and Supplementary Figure 1).

**Figure 4.**
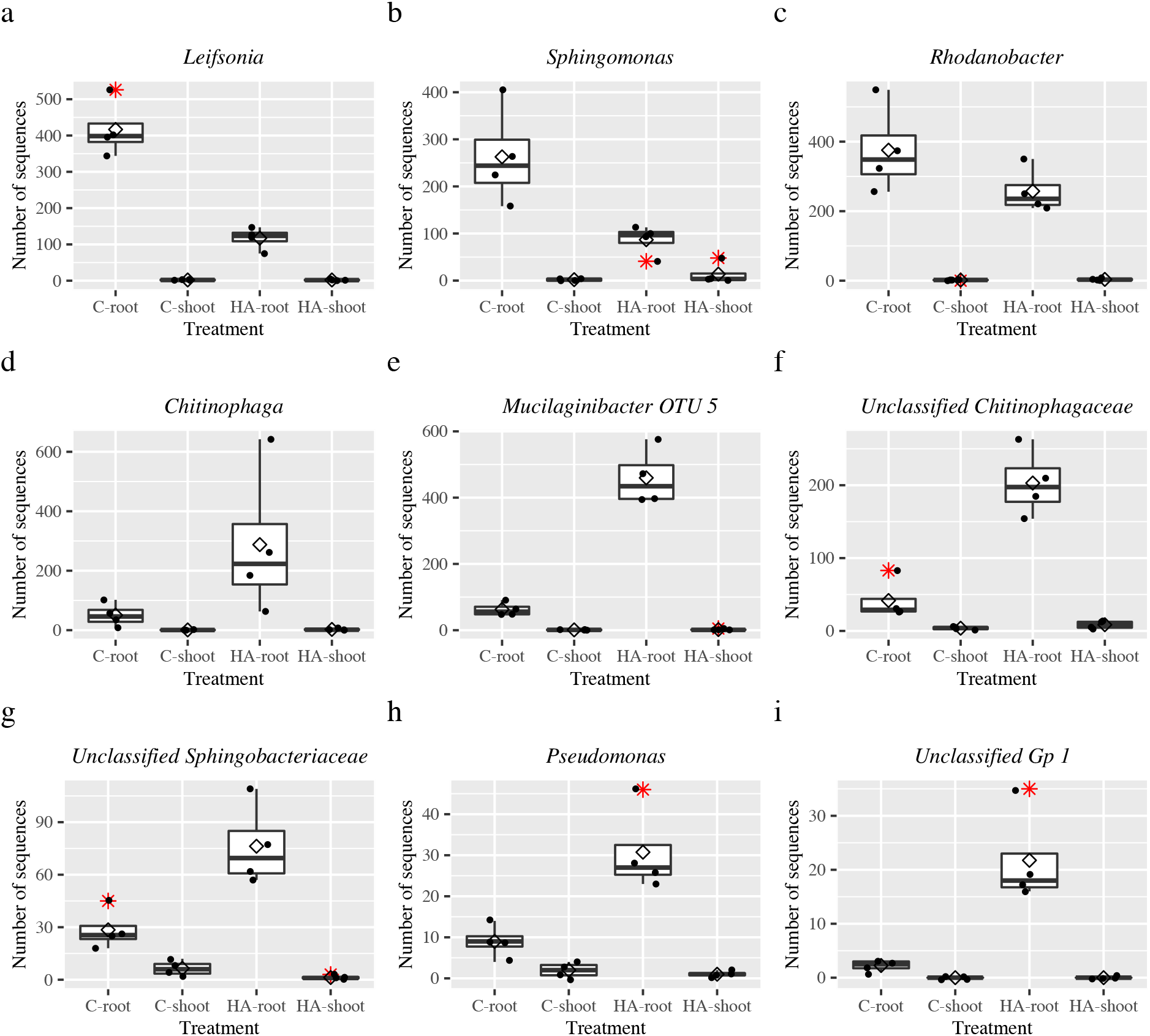
Major taxa whose abundance were differently decreased (a, b, and c) or enriched (d through i) in roots that received humic acids (HA). C- shoot, C- shoot, HA-shoot, and HA- shoot stand for shoot and roots of the control and humic-acid-treated plants, respectively. The abundances are represented by the number of sequences. The white diamond identifies the average count number, and the asterisks represent outliers. Differential abundances were detected as significant by the differential gene expression analysis on the negative binomial distribution implemented in package DESeq2 (Love et al., 2014).

## Discussion

In this work, we evaluated the effect of HA on the growth and root morphology of rice plants together with the identification of bacterial taxa selected under this condition. We believe there is a synergism between HA and the plant-associated microbiota benefiting the plant since both can modulate plant physiological processes (Hallmann and Berg, 2006; Canellas and Olivares, 2014). HA beneficial effects could derive from its direct action on the plant metabolism or through the stimulation of the plant microbiota.

We observed higher growth rates (figure 1) in HA-treated plants. This increase was related to higher biomass production efficiency, indicating an increase in photosynthetic efficiency and assimilation of nutrients. In line with our data, additional studies reported that HA stimulated photosynthesis (Nardi *et al*., 2002; Jannin *et al*., 2012). There are also reports on the ability of HS to stimulate different components of the primary plant metabolism, including the activity of chlorophylls a and b (Yang *et al*., 2004) and different components of carbon metabolism, mainly those related to the stimulation of enzymes (Nardi *et al*., 2007), carbohydrate production (Merlo *et al*., 1991), and expression of energy metabolism genes (Trevisan *et al*., 2011). For example, Jannin *et al.* (2012) showed that the application of HA in rapeseed roots provided a significant improvement in the liquid photosynthetic rate, the number of chloroplasts per cell, and starch granules. These effects were consistent with the increase C, N, and S assimilations. These authors also documented HA induced root growth, a process that was also observed in our work. Using a temporal evaluation approach, we observed that the increments on root and plant biomass varied with the moment of observation (Figures 1 and 2). There are not many reports of experiments that evaluate these parameters in a daily sequence; however, the stimulating effect of HA on root growth and development has been reported by several authors such as Canellas *et al*. (2002); Zandonadi *et al*. (2010); Mora *et al*. (2012); and Garcia *et al*. (2016*b*).

The metataxonomics analysis showed that the plant-associated bacterial community responded to the presence of HA. In hydroponic conditions, the shoot showed higher bacterial diversity than the roots in both treatments. The presence of HA led to prominent variations in root community composition and, to a lesser extent, in the shoot (Figure 3). Among the main variations are the increased *Bacteroidetes* and *Acidobacteria* increased in the roots, and the increased abundance of *Proteobacteria* with a decrease of *Bacteroidetes* in the shoots. *Rhizobium*, a proteobacterium and a genus commonly associated with legumes (Long, 2001), was the most abundant genus, especially in the roots. Sturz *et al*. (1997) reported that the presence of this genus in clover, a legume, was not restricted to the nodules, but that it could be found in the whole plant. These authors also found higher bacterial diversity in the leaves a result similar to the one we observed here. This genus is capable of colonizing rice (Yanni *et al*., 1997; Hahn *et al*., 2016; Rosenblueth *et al*., 2018) and has been reported as promoting rice growth when inoculated alone or in mixture with *Azospirillum brasilense* (Hahn *et al*., 2016).

The genera with the highest abundance in the roots upon the application of HA were *Chitinophaga* and *Mucilaginibacter* (Figure 3c, and Figure 4d,e). Both genera have members that produce a range of hydrolytic enzymes, e.g., glucanases and chitinases, involved in the degradation of complex polysaccharides, such as those in the cell wall of fungi, oomycetes, and nematode eggs (Ramamoorthy *et al*., 2001, Yoon *et al*., 2012, Mélida *et al*., 2013; Carrion *et al*., 2019; Sharma *et al*., 2020). These same genera were identified in rhizosphere soils after applying a compound that induces systemic resistance of plants to diseases (Nicot *et al*., 2016) and in association with plants resistant to pathogens (Dai *et al*., 2020). Besides, the invasion of sugar beet roots by *Rhizoctonia solani* stimulated the enrichment of members of *Chitinophagaceae* in plants cultivated in disease-suppressive soil (Carrion *et al*., 2019). Members of this group contributed to the pool of debranching enzymes and enzymes associated with fungal cell-wall degradation. Together with others (e.g., siderophore production), these functional characteristics could provide plant protection from the inside out. *Pseudomonas and Gp 1* were also stimulated HA-treated roots, although their abundances were ten to twenty times lower than the abundance of *Chitinophaga* and *Mucilaginibacter. Pseudomonas* is widely known as a plant growth-promoting bacterium, and it is commonly associated with the control of phytopathogens (Ramamoorthy *et al*., 2001). In addition to the direct action of inducing the plant defense, bacteria from this genus synthesize a variety of antimicrobial compounds and hydrolytic enzymes capable of degrading components of the fungal cell wall (Ramamoorthy *et al*., 2001; Wang *et al*., 2019). *Gp1* has been recently described as promoting plant growth through mechanisms such as idol-3-acetic acid and siderophore production (Kielak *et al*., 2016). We highlight that this second mechanism has also been reported as a mechanism involved in the inhibition of plant pathogens (Koepler *et al*., 1980). Altogether, these characteristics call attention to the possible role of the bacterial community selected in HA-treated plants on the protection of the plant against pathogens.

Recent studies have shown that the rhizosphere community of disease-suppressing soil acts as the first line of plant defense (Mendes *et al*., 2011). When this barrier is broken, pathogens face defense mechanisms induced by the plant (Jones and Dangl, 2006). Following the invasion and entering the plant tissue, the microbiota’s modulation could act as another protective layer, the third line of defense against the progress of the infection (Carrión *et al*., 2019). This last study found both *Pseudomonadaceae* and *Chitinophagaceae* are enriched in sugar beet grown in a pathogen-inoculated suppressive soil. Especially members of the second family contributed to with debranching enzymes and enzymes associated with fungal cell-wall degradation. It is noteworthy that an increase in similar groups was achieved here upon the application of humic acids.

For the first time, we call attention to the possible role of HAs to stimulate members of the plant-associated bacterial community that may protect the plant against pathogens. Given these results, we suggest that plants recruit these microorganisms in response to the stress caused by the HA-root interaction (Berbara and Garcia, 2014; García *et al*., 2016a). When HAs are applied on plant roots, as occurred here, HA fragments agglomerate on the root surface and cause mild and transient stress. In response, the plants modulate signaling through reactive oxygen species (ROS), inducing a physiological state of protection to abiotic stress (Berbara and Garcia, 2014; García *et al*., 2016a). This stress may be one of the mechanisms stimulating shifts in the composition of communities and the enrichment of bacteria potentially involved in plant defense. Previous studies also showed that the co-application of bacteria and HA improved the recovery of bean plants from water stress, further supporting our hypothesis (Olivares *et al*., 2017). These same authors reported different sugarcane responses when exposed to bacteria, HAs, or their mixture under water restriction. The activities of antioxidant enzymes such as superoxide dismutase, catalase, and ascorbate peroxidase remained higher after rehydration only in HA-treated plants.

Studies also point out to the role of humic substances on plant resistance to biological agents and the effect of these substances on secondary plant defense metabolism (Polak and Pospíšil, 1995; Abdel-monaim et al., 2011). HA-treated plants showed higher expression of phenylalanine (tyrosine) ammonia-lyase (PAL / TAL; EC 4.3.1.5), an enzyme that catalyzes the first step in the biosynthesis of phenolic compounds (Schiavon *et al*., 2010; Canellas *et al.,* 2014). These are secondary metabolites that protect plants against a variety of biotic and abiotic stresses (Dixon and Paiva, 1995). It has been suggested that the effect of HAs on the metabolism of phenylpropanoids is associated with their chemical structure, co-purified fungal elicitors, and other signaling molecules of these substances (Schiavon *et al.*, 2010).

## Conclusions

We can conclude that the application of HAs in plants modifies the composition of the bacterial community of rice simultaneously with the stimulation of root growth and development. We also suggest that HAs can trigger the enrichment of microorganisms that act on the defense of the plant against pathogens. These results are unprecedented in the literature and evince previously open questions, such as the need for approaches to elucidate the real role of HA on plant physiology and the participation of the plant microbiota. Once confirmed, our findings may point out to the use of HA as part of a strategy to prime plants against biotic and abiotic stresses through the stimulation of their defense metabolism, including the recruitment of pathogen-suppressing microorganisms.

## Acknowledgments

We thank the Carlos Chagas Filho Foundation for the Support of Research in the State of Rio de Janeiro (FAPERJ) for its financial support (project E-26/202.683/2018); the Brazilian National Council for Scientific and Technological Development (CNPq) for the research fellowship provided to Ederson da Conceição Jesus (project 475168/2012-7); and the Coordenação de Aperfeiçoamento de Pessoal de Nível Superior (CAPES) for the scholarship provided to Maura Santos Reis de Andrade da Silva. This work is also sponsored by the INCT - Plant-Growth Promoting Microorganisms for Agricultural Sustainability and Environmental Responsibility (CNPq, 465133/2014-4, Fundação Araucária-STI, CAPES). We thank Sarah Owens and Stephanie M. Greenwald, both from the Argonne National Laboratory, for supporting us with the sequencing analysis.

## Authors contributions

M.S.R.A.S. designed the experiments with advice from O.C.H.T., A.C.G., J.G.M., and R.L.L.B., who advised on the HA-application assay, and E.C.J, and V.L.D.B. who advised about the microbiological analyses. The experiment was carried out with the assistance of O.C.H.T., C.S.R.A.S., and C.S.R.A.S. Data analysis was performed with support from O.C.H.T., T.G.R., and E.C.J. M.S.R.A.S. wrote the paper. All coauthors revised it and contributed to its final version.

